# Fewer non-native insects in freshwater than in terrestrial habitats across continents

**DOI:** 10.1101/2022.03.22.485042

**Authors:** Agnieszka Sendek, Marco Baity-Jesi, Florian Altermatt, Martin K.-F. Bader, Andrew M. Liebhold, Rebecca Turner, Alain Roques, Hanno Seebens, Piet Spaak, Christoph Vorburger, Eckehard G. Brockerhoff

## Abstract

**Aim:** Biological invasions are a major threat to biodiversity in both aquatic and terrestrial habitats. Insects represent an important group of species in freshwater and terrestrial habitats, and they constitute a large proportion of non-native species. However, while many non-native insects are known from terrestrial ecosystems, it remains unclear how they are represented in freshwater habitats. Comparisons of the richness of invaders relative to the richness of native species between freshwater and terrestrial habitats are scarce, which hinders syntheses of invasion processes. Here, we used data from three regions on different continents to determine whether non-native insects are under- or overrepresented in freshwater compared to terrestrial assemblages.

**Location:** Europe, North America, New Zealand

**Methods:** We compiled a comprehensive inventory of the native and non-native insect species established in freshwater and terrestrial habitats of the three study regions. We then contrasted the richness of non-native and native species among freshwater and terrestrial insects for all insect orders in each region. Using binomial regression, we analysed the proportions of non-native species in freshwater and terrestrial habitats across the three regions.

**Results:** In most insect orders living in freshwater, non-native species were under-represented, while they were over-represented in a number of terrestrial orders. This pattern occurred in purely aquatic orders as well as in orders with both freshwater and terrestrial species. Overall, the proportion of non-native species was significantly lower in freshwater than in terrestrial species.

**Main conclusions:** Despite the numerical and ecological importance of insects among all non-native species, non-native insect species are surprisingly rare in freshwater habitats, and this pattern is consistent across the three investigated study regions. We briefly review hypotheses concerning species traits and invasion pathways that are most likely to explain these patterns. Our findings contribute to a growing appreciation of drivers and impacts of biological invasions.

## 1. Introduction

Biological invasions are an important component of global change (IPBES, 2019; Pyšek et al., 2020) and a well-recognized threat to biodiversity in both terrestrial (D’Antonio & Kark, 2002) and aquatic habitats (Francis & Chadwick, 2021; Jackson et al., 2017). Freshwater ecosystems are not only highly invaded (Bolpagni, 2021; Pyšek et al., 2010) but also very susceptible to severe impacts of invasive species (Moorhouse & Macdonald, 2015; Sala et al., 2000). Macroinvertebrates are common and often damaging invaders of freshwater ecosystems worldwide (Baur & Schmidlin, 2007; Cuthbert et al., 2021; Emery-Butcher et al., 2020; Ricciardi, 2015). However, the representation of different higher-level taxa among non-native aquatic invertebrates is unbalanced (Ricciardi, 2015). For example, aquatic insects have fewer non-native representatives than other groups of aquatic invertebrates such as crustaceans and molluscs (Fenoglio et al., 2016; Karatayev et al., 2009; Ricciardi, 2015).

This imbalance is surprising as insects are a highly species rich, diverse and ubiquitous taxon (Chapman, 2009; Stork, 2018) occurring in nearly all terrestrial and freshwater habitats. Furthermore, non-native insects vastly outnumber other invertebrates and vertebrates in species richness (Roques et al., 2010; Seebens et al., 2017). However, Liebhold et al. (2016) suggested that insect orders that are dominated by aquatic species (*i*.*e*., Ephemeroptera, Odonata, Plecoptera or Trichoptera) have relatively few non-native representatives. Yet, it is unclear whether this pattern also occurs among insect orders comprising both terrestrial and aquatic species. Moreover, to our knowledge, the question whether aquatic insects are indeed less common invaders than terrestrial insects has not been explicitly investigated on a large taxonomic and geographic scale.

The number of established non-native species is increasing worldwide (Seebens et al., 2017), and so is the colonisation by aquatic and terrestrial invertebrates (Baur & Schmidlin, 2007; Brockerhoff & Liebhold, 2017; Ricciardi, 2006). Exploring patterns of invasions across habitats can advance our understanding of invasion processes and drivers and provide crucial information informing efforts to mitigate future invasions.

Here, we examined the proportions of established non-native freshwater and terrestrial insects relative to the number of corresponding native species across three major geographic regions. In particular, we analysed whether freshwater non-native insects are indeed less likely to invade than terrestrial insects and whether this is a universal phenomenon across all insect orders and across continental regions (Europe, North America and Australasia *i*.*e*., New Zealand). Finally, we elaborated on these results with regard to hypotheses related to life history traits, habitat specifics, and invasion pathways which may explain such differences between freshwater and terrestrial insects.

## 2. Methods

### 2.1 Data compilation

We collected data on the numbers of native and non-native insect species. In compiling lists of non-native species, we considered established non-native species with self-sustaining populations outdoors (Colautti & MacIsaac, 2004; Pyšek et al., 2004). Our datasets did not include non-native species which, to our knowledge, occur only in indoor conditions including aquaria and greenhouses. Species lists were collected for freshwater and terrestrial habitats in Europe, North America, and New Zealand, respectively, for which comprehensive lists of native and non-native insect species are available. The information was assembled from multiple published species inventories (details provided in the Note S1; Foottit *et al*., 2017; de Jong *et al*., 2014; Liebhold et al., 2016; Simpson et al., 2018; Turner et al. 2021a; Yamanaka *et al*., 2015, recently updated in an online database: Turner et al. 2021b), books (Macfarlane *et al*., 2010; Merritt & Cummins, 1996; Merritt *et al*., 2008) and online databases (DAISIE and EASIN databases; Roques *et al*., 2010; 2016) and freshwaterecology.info (Schmidt-Kloiber & Hering 2015). We standardised taxonomic information according to the GBIF Backbone Taxonomy (Secretariat, G.B.I.F. 2021), with the exception of the order Psocodea which we kept as two separate orders, Phthiraptera and Psocoptera, as they have been treated traditionally. In this way, we obtained 29 orders with at least one species, either native or non-native (Table S1; Table S2).

There is no single rigorous definition of what constitutes an aquatic *vs*. a terrestrial insect species. In fact, definitions of aquatic and terrestrial insects differ greatly between studies (compare Merritt & Cummins 1996; Merritt et al. 2008). Here, we applied a strict and a liberal definition of freshwater aquatic species (while treating all others as “terrestrial”). In the “strict” definition, we consider as freshwater insects those species which spend at least one stage of their life cycle obligatorily in freshwater environments. This definition encompasses insects living on the surface of lotic and lentic water bodies, including temporal habitats such as puddles or water-filled tree holes. In the “liberal” definition, we also included insects occupying semi-aquatic habitats such as the edges of water bodies, or algal mats, species specialised on feeding on hydrophytes (including burrowers and miners) in submergent, emergent and floating zones, or living in water-saturated substrates such as wet soil or wood, or sap flows. For the liberal dataset we also included parasitoids known or suspected to attack hosts under the water surface (Burghele 1959; Merritt & Cummins 1996). In both of these classification systems, insects feeding on terrestrial vegetation growing in proximity to water bodies and parasitoids of terrestrial stages of aquatic insects were not considered aquatic (Merritt & Cummins 1996; Merritt et al. 2008).

### 2.2 Data analysis and presentation

To compare numbers of non-native and native insect species in freshwater and terrestrial habitats, we computed the total number of native and non-native species, in both kinds of habitat, for each of the three regions and insect order. The overall ratio of non-native to native freshwater and terrestrial species were compared using Chi-square tests for each of the three regions separately. Then, we compiled these values by insect order. For this, we overlaid a plot showing the number of freshwater and terrestrial species in each order with a line representing the number of non-native species per order that would be expected if they were in the same proportions as all the non-native species to all the native species in the region. We calculated a 95 % prediction interval based on a binomial distribution under the assumption that the proportions of non-native and native species in a given order are identical to the overall proportions in each of the regional datasets. Insect orders that were located above or below the line and outside of the confidence intervals were considered over- or under-represented, respectively, in terms of the number of non-native species.

We compared non-native to native species ratios in freshwater and terrestrial habitats by using a generalized mixed effect regression model with binomial error distribution and a probit link function. Insect order was accounted for by including it as a random term. As fixed effects, we used regions (Europe, North America, New Zealand), habitat (freshwater, terrestrial) and their interaction. We applied a single term deletion based on likelihood ratio testing to remove non-significant terms. To conduct pairwise comparisons of habitats across the regions, we estimated and contrasted marginal means (Kaltenbach, 2021) with a Benjmaini & Hochberg correction for multiplicity adjustment (Benjamini & Hochberg, 1995). All analyses were done in R version 4.0.5 (R Core Team, 2021). We computed regression models using the `glmmTMB` package (Brooks et al., 2017; Magnusson et al. 2017) and used the `DHARMa` package (Hartig, 2021) to examine the residuals. Pairwise comparisons of marginal means were performed using the `emmeans` package (Lenth, 2021).

## 3. Results

### 3.1 All freshwater vs. terrestrial insects

Overall, freshwater insects are much less species rich than terrestrial insects, in both native and non-native species (Fig. 1; Fig. S1; Fig. S2). Moreover, the numbers of non-native species were considerably lower relative to the numbers of native species across all regions, which was reflected in the small ratios of non-native to native species numbers (almost 10-fold in Europe and North America and almost 4-fold in New Zealand; Table 1; Fig. S2). The results were similar using the liberal dataset (Table 1; Fig. S2).

**Table 1.**
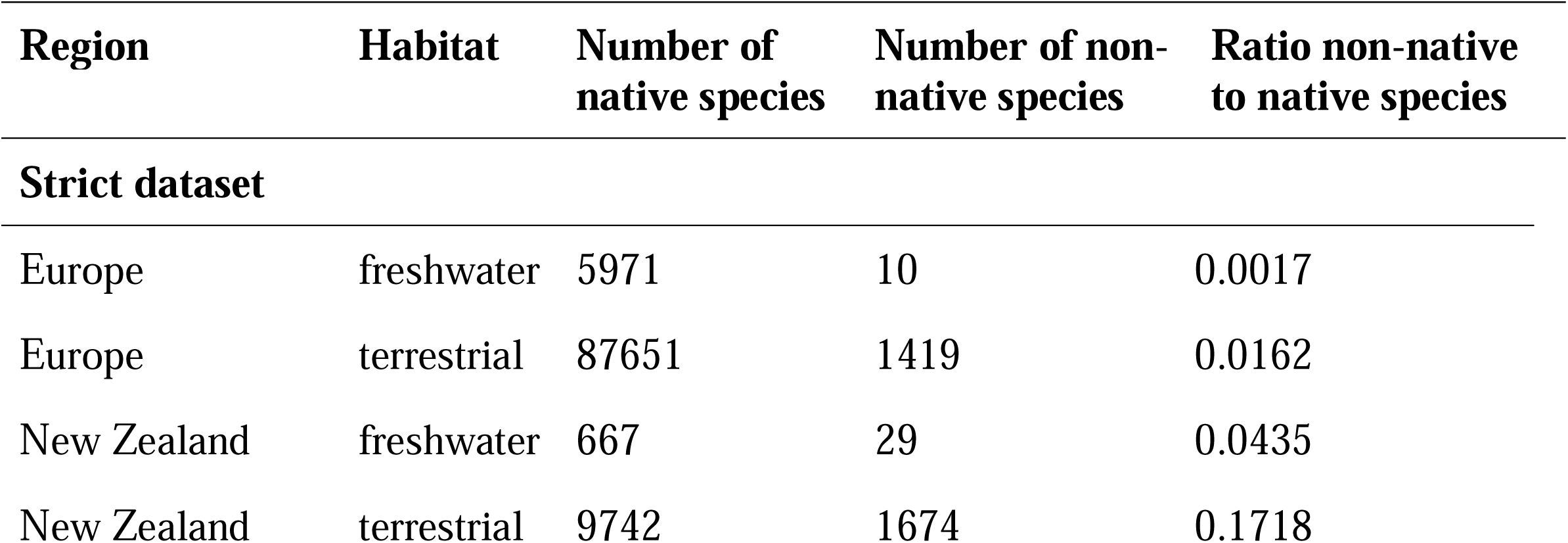
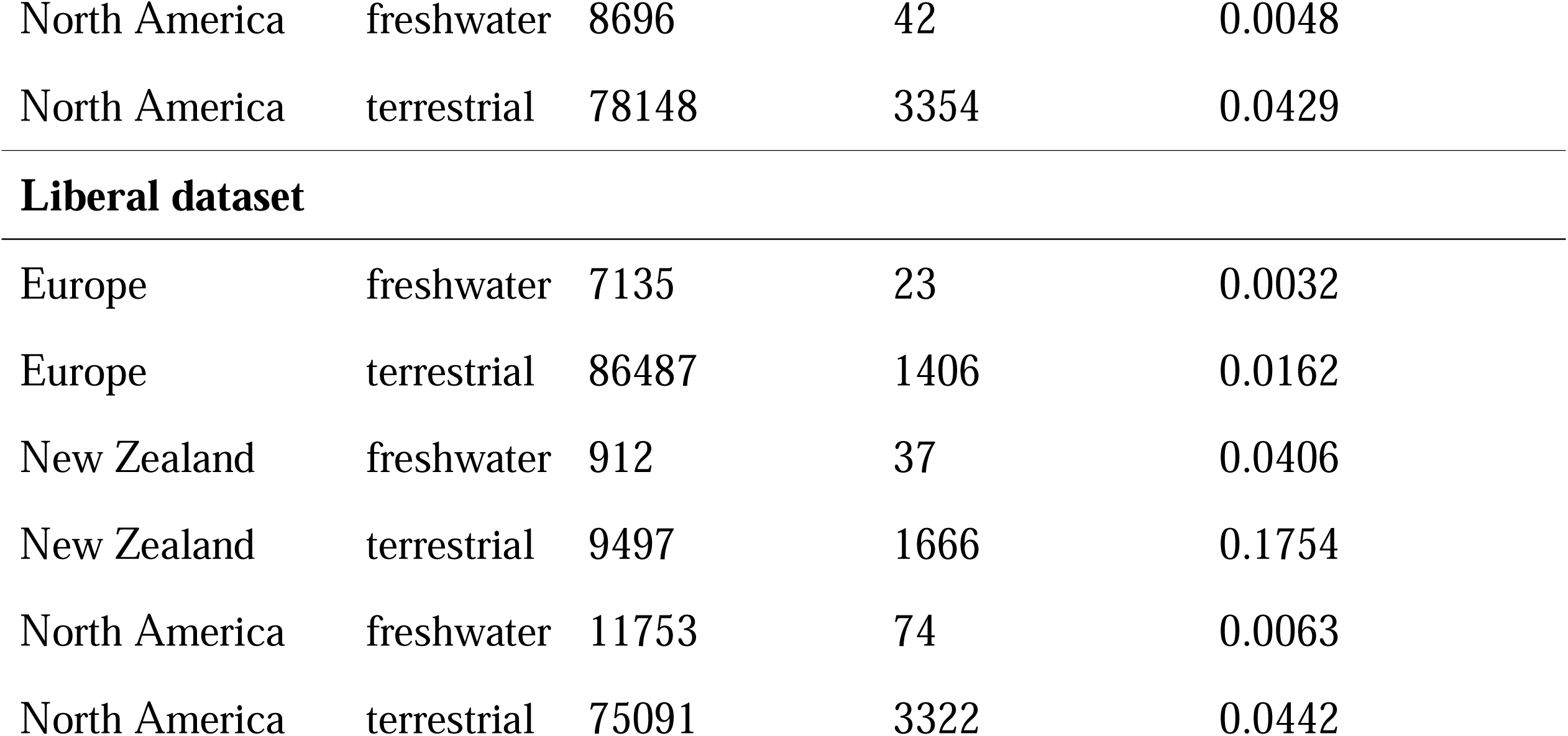
Overall number of freshwater and terrestrial native and non-native insects in Europe, New Zealand, and North America) and ratios of non-native to native species.

| Region | Habitat | Number of native species | Number of non-native species | Ratio non-native to native species |
| --- | --- | --- | --- | --- |
| Strict dataset |  |  |  |  |
| Europe | freshwater | 5971 | 10 | 0.0017 |
| Europe | terrestrial | 87651 | 1419 | 0.0162 |
| New Zealand | freshwater | 667 | 29 | 0.0435 |
| New Zealand | terrestrial | 9742 | 1674 | 0.1718 |
| North America | freshwater | 8696 | 42 | 0.0048 |
| North America | terrestrial | 78148 | 3354 | 0.0429 |
| **Liberal dataset** |  |  |  |  |
| Europe | freshwater | 7135 | 23 | 0.0032 |
| Europe | terrestrial | 86487 | 1406 | 0.0162 |
| New Zealand | freshwater | 912 | 37 | 0.0406 |
| New Zealand | terrestrial | 9497 | 1666 | 0.1754 |
| North America | freshwater | 11753 | 74 | 0.0063 |
| North America | terrestrial | 75091 | 3322 | 0.0442 |

**Fig. 1.**
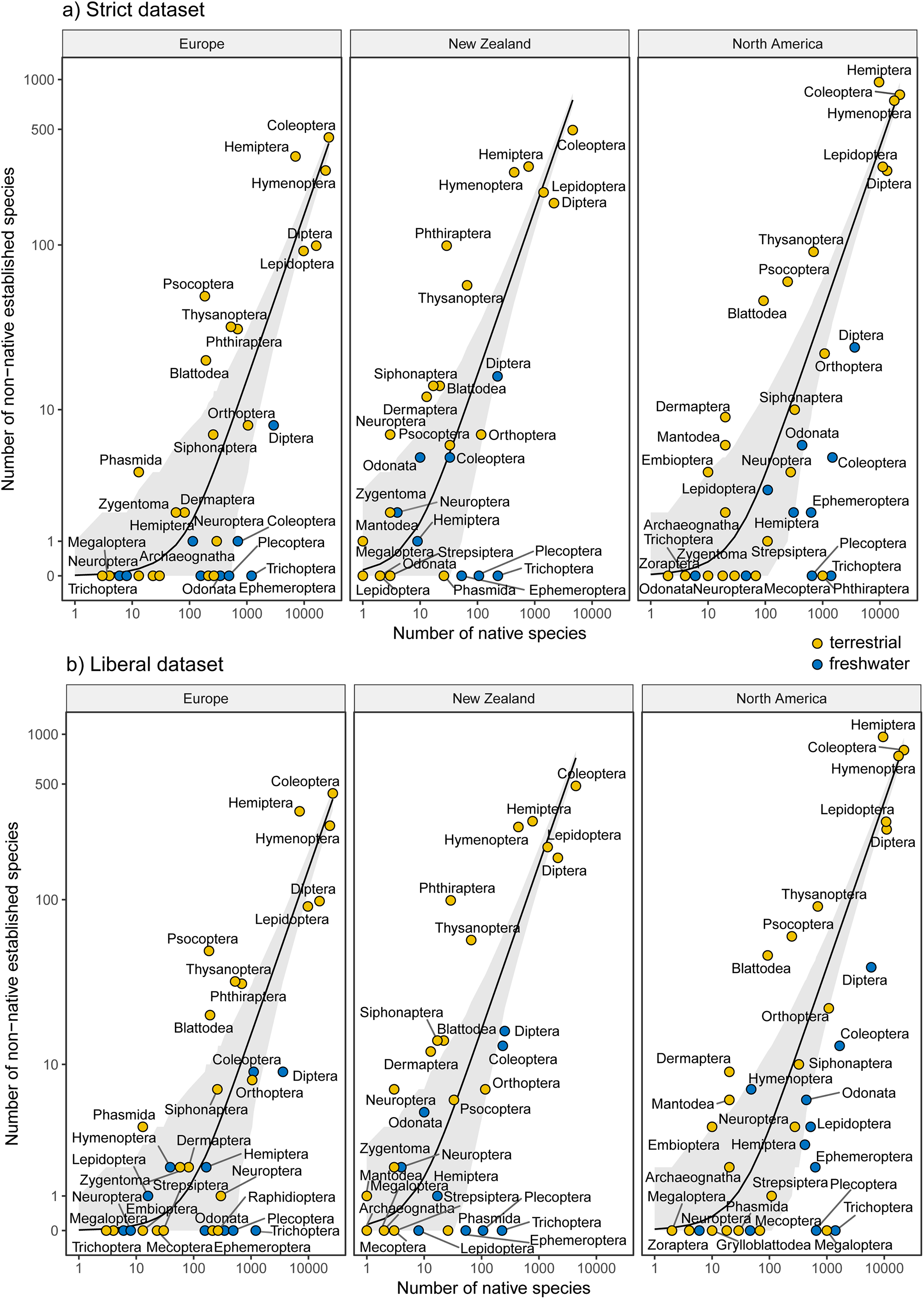
Numbers of native versus non-native species per insect order in freshwater (blue) and terrestrial (yellow) habitats across the three investigated regions (Europe, New Zealand, and North America). Solid lines represent the expected number of non-native species assuming that the proportions of non-native and native species in a given order are identical to the overall proportions in each of the regional datasets. The shading represents the 95% confidence interval, based on the binomial distribution. Orders located outside of the shaded range are considered under-or overrepresented. Panels (a) and (b) represent strict and liberal datasets, respectively.

### 3.2 Comparison at the level of insect orders

At the order level, we found that many of the freshwater insects were significantly under-represented in terms of their proportions of non-native species (*i*.*e*., they fell below the expectation line and outside the 95 % prediction interval, Fig. 1). Among them were orders that are purely or almost purely aquatic (*e*.*g*., Ephemeroptera, Plecoptera, Trichoptera) as well as aquatic subsets of the larger orders that contain both freshwater and terrestrial species (*e*.*g*., Coleoptera and Diptera). By contrast, non-native species were over-represented in numerous purely terrestrial insect orders (*e*.*g*. Blattodea, Hemiptera, Phthiraptera, Psocoptera, Thysanoptera) (*i*.*e*., they fell above the expectation line and outside the 95 % prediction interval, Fig. 1). These patterns were broadly similar across the three regions. Likewise, the results were comparable for the strict and liberal datasets, although some groups (*e*.*g*., aquatic Coleoptera) shifted slightly (Fig. 1).

The mean proportions across insect orders of non-native species (out of all species) were significantly lower for freshwater insects than for terrestrial insects, and this was consistent across the three regions (Table 2; Fig. 2; Table S3; Table S4). The magnitude of differences between regions varied between the strict and the liberal datasets (Table 2; Fig. 2). The difference between non-native freshwater and terrestrial species was only marginally non-significant for the strict dataset fromNew Zealand, while there was a significant difference for the liberal dataset (Table 2; Fig. 2).

**Table 2.**
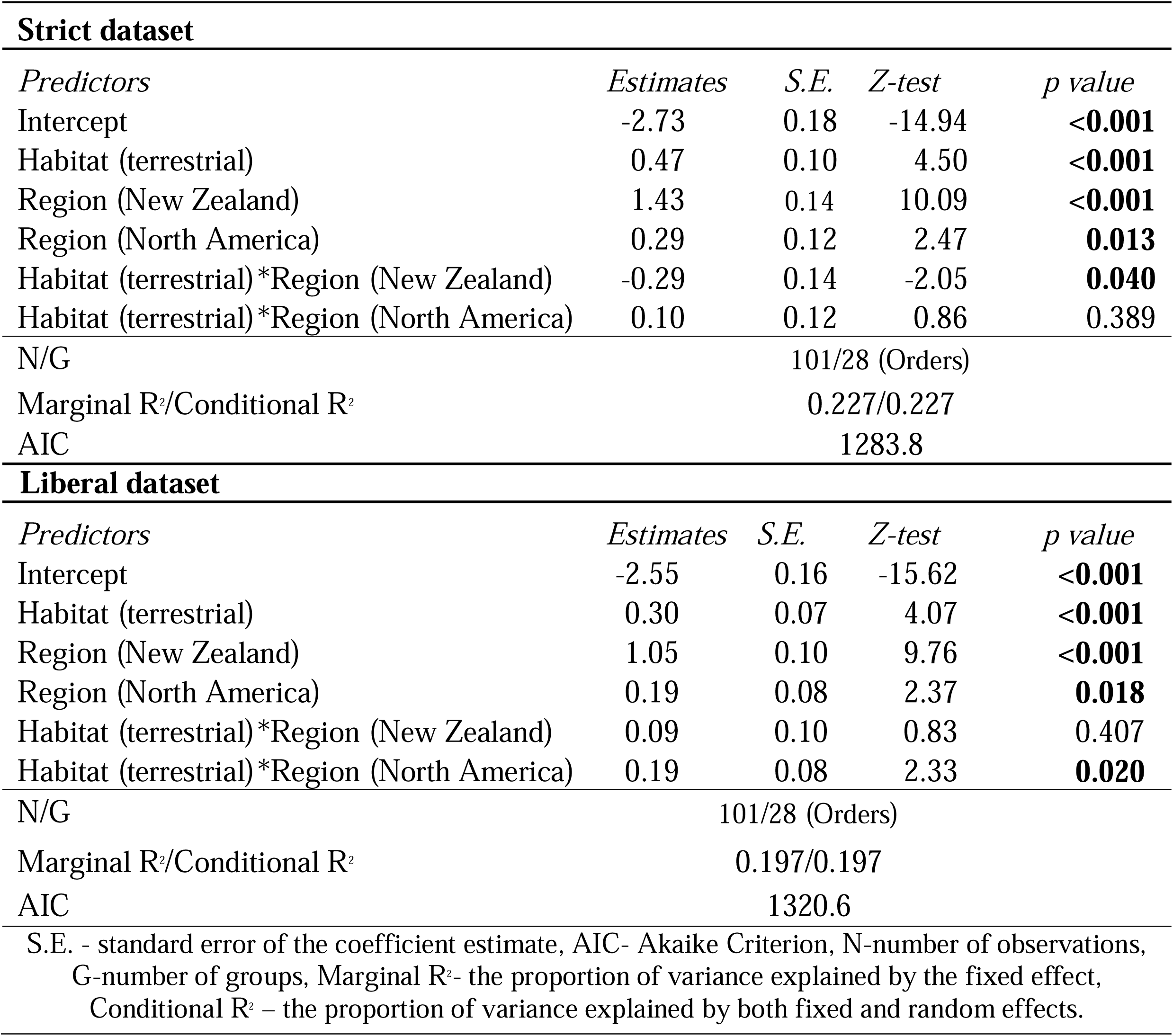
Summary of effects of habitat (freshwater versus terrestrial) and region (Europe, New Zealand, and North America) on the ratio of non-native to native species in the insect orders. The table represents the outcome of the binomial regression analysis of the strict and liberal datasets. In this analysis, the data from Europe were used as the reference point so that the estimates shown are relative to the results for Europe.

| Strict dataset |  |  |  |  |
| --- | --- | --- | --- | --- |
| Predictors | Estimates | S.E. | Z-test | p value |
| Intercept | -2.73 | 0.18 | -14.94 | <0.001 |
| Habitat (terrestrial) | 0.47 | 0.10 | 4.50 | <0.001 |
| Region (New Zealand) | 1.43 | 0.14 | 10.09 | <0.001 |
| Region (North America) | 0.29 | 0.12 | 2.47 | 0.013 |
| Habitat (terrestrial)*Region (New Zealand) | -0.29 | 0.14 | -2.05 | 0.040 |
| Habitat (terrestrial)*Region (North America) | 0.10 | 0.12 | 0.86 | 0.389 |
| N/G |  | 101/28 (Orders) |  |  |
| Marginal R²/Conditional R² |  | 0.227/0.227 |  |  |
| AIC |  | 1283.8 |  |  |
| Liberal dataset |  |  |  |  |
| Predictors | Estimates | S.E. | Z-test | p value |
| Intercept | -2.55 | 0.16 | -15.62 | <0.001 |
| Habitat (terrestrial) | 0.30 | 0.07 | 4.07 | <0.001 |
| Region (New Zealand) | 1.05 | 0.10 | 9.76 | <0.001 |
| Region (North America) | 0.19 | 0.08 | 2.37 | 0.018 |
| Habitat (terrestrial)*Region (New Zealand) | 0.09 | 0.10 | 0.83 | 0.407 |
| Habitat (terrestrial)*Region (North America) | 0.19 | 0.08 | 2.33 | 0.020 |
| N/G |  | 101/28 (Orders) |  |  |
| Marginal R²/Conditional R² |  | 0.197/0.197 |  |  |
| AIC |  | 1320.6 |  |  |
| S.E. - standard error of the coefficient estimate, AIC- Akaike Criterion, N-number of observations, G-number of groups, Marginal R²- the proportion of variance explained by the fixed effect, Conditional R² – the proportion of variance explained by both fixed and random effects. |  |  |  |  |

**Fig. 2.**
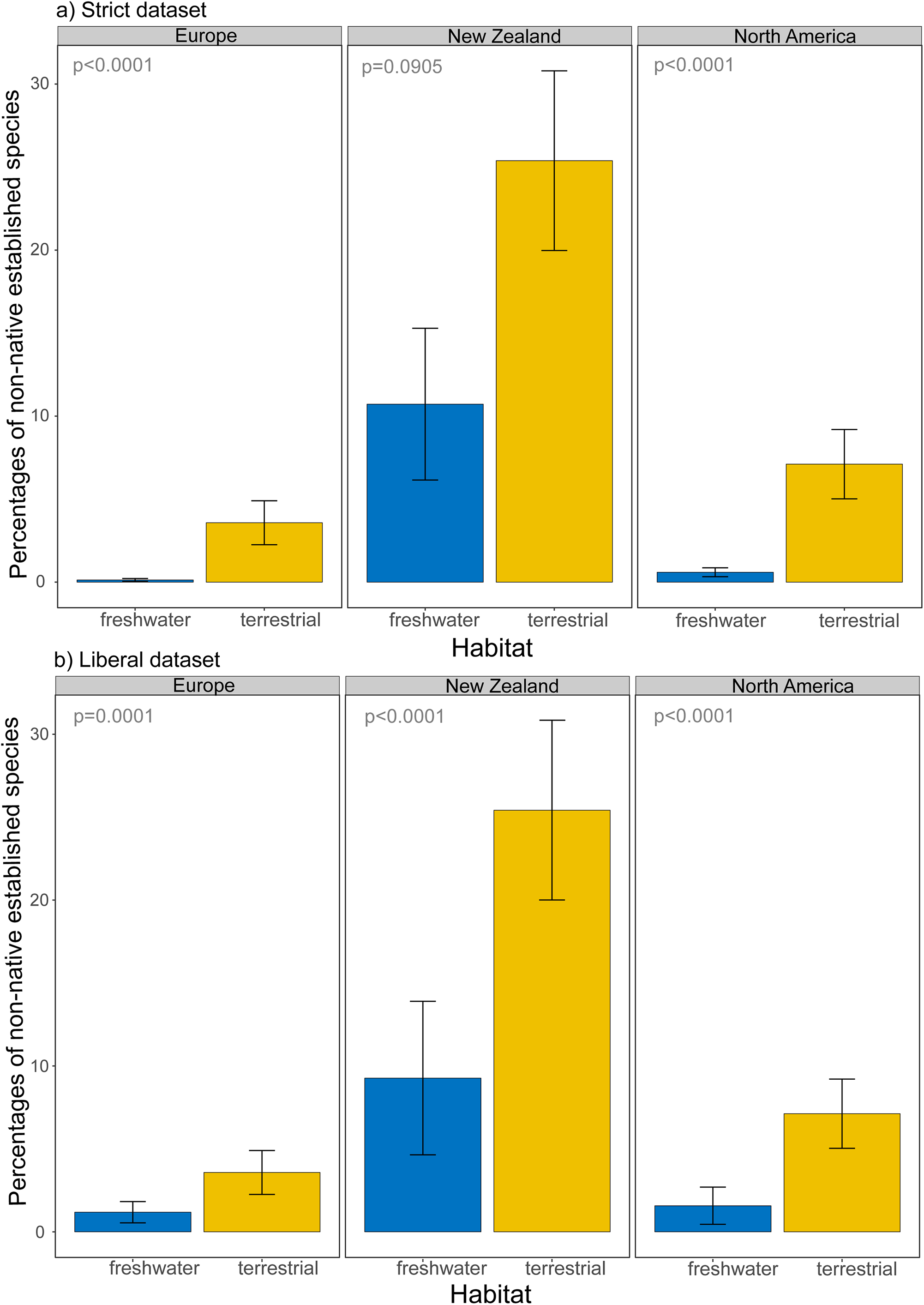
Percentages (calculated from proportions) of non-native, established species (out of all species) in freshwater and terrestrial habitats across the three investigated regions (Europe, New Zealand, North America). The error bars represent the standard error around the mean. *P*-values indicating differences between the proportions of freshwater and terrestrial species within regions are obtained from marginal means contrasts. Panels (a) and (b) represent datasets based on strict and liberal definitions of aquatic insects, respectively.

### 3.2 Taxa most represented among invaders

Diptera stand out as the most species-rich group of non-native freshwater insects in all three regions, whereas Coleoptera, Hemiptera and Hymenoptera dominate in terrestrial habitats (Fig. 1; Table S2). Prominent dipteran families among non-native freshwater insects are the Culicidae (mosquitoes), Chironomidae (non-biting midges) and Syrphidae (hover flies). Only three species (all dipterans) occurred in more than one region as non-native species: the Asian tiger mosquito (*Aedes albopictus*), the southern house mosquito (*Culex quinquefasciatus*), and the common drone fly (*Eristalis tenax*) (Table S2). Despite the prominence of non-native freshwater Diptera, they are consistently under-represented relative to the number of native species (Fig. 1). Even though Odonata are a comparatively small order, they are the second-most common order of freshwater invaders in New Zealand and North America (although they are absent in Europe) (Fig. 1, Table S2). Of the other typical freshwater insects, only two species of mayflies (Ephemeroptera) were among the non-native species, both in North America.

## 4. Discussion

Our analyses show that freshwater insects are consistently under-represented compared to terrestrial species in terms of their numbers of non-native species relative to native species. This was evident in the non-native to native species ratios which were considerably lower in freshwater than in terrestrial insects. The pattern of fewer non-native species in aquatic insects than in terrestrial insects occurred in purely aquatic orders as well as in orders which contain both freshwater and terrestrial representatives. The effect was consistent across the three study regions (Europe, North America and New Zealand), which indicates that this under-representation of non-native freshwater insects is occurring at a large geographical scale, and may be inherent and generalizable across insects overall. Our findings are remarkable because insects (in general) are the most species-rich class (Chapman, 2009; Stork, 2018), and they are more numerous as established non-native species than any other animal group (Seebens et al., 2017). However, despite the high richness of aquatic insects (Balian et al., 2008; Dijkstra et al., 2014) and their high abundance in all kinds of freshwater habitats (Hershey et al. 2001), there are surprisingly few freshwater insect species that have invaded non-native regions.

An apparent paucity of non-native freshwater insects has been noted previously, mainly in comparison with other groups of macroinvertebrates, such as Crustacea or Mollusca (Fenoglio et al., 2016; Ricciardi, 2015), rather than in comparison with terrestrial insects. Other studies have concluded that there are very few invaders among freshwater insects despite their high species richness in the native benthic fauna (Karatayev et al., 2009; Ricciardi, 2015). Our study is the first to demonstrate conclusively that freshwater insects *per se* are less common as invaders compared with terrestrial insects. This is relevant because there are very few comparisons between freshwater and terrestrial habitats in terms of the richness of invaders relative to the richness of native species (though see Liebhold et al., 2016). Furthermore, our analyses, based on comprehensive inventories of insects from three well-studied large regions far from each other and with different control processes at borders (Europe, North America and New Zealand), suggest that this under-representation of non-native freshwater insects is a universal pattern.

### Drivers and mechanisms

The mechanisms responsible for the limited success of freshwater insects as invaders are not fully understood. However, based on theoretical considerations and the characteristics of the few freshwater insect species that succeeded to invade, a number of hypotheses can be discussed.

### Transport pathways and interactions

Transport pathways typically involved in invasions of aquatic species and the relative unsuitability of these pathways for freshwater insects (compared with other freshwater species) are likely to play a key role. Several recent studies highlighted the importance of pathways in insect invasions (Liebhold et al., 2012; 2016; Ricciardi, 2006; Turner et al., 2021a). While numerous terrestrial non-native insects are being transported with their host plants via trade in plant products and live plants (Brockerhoff & Liebhold, 2017; Liebhold et al., 2012), this is thought to be a less likely pathway for aquatic insects (Fenoglio et al., 2016), the latter being generally much less associated with specific host plants. For example, numerous species of terrestrial Thysanoptera, an over-represented terrestrial order (see Fig. 1; Fig. S1) are transported with their crop or ornamental host-plants (Liebhold et al., 2016). By contrast, we observed only a single representative of a semi-aquatic Thysanoptera (*Organothrips indicus*) (in Europe). However, this species, along with nine herbivorous Crambidae (Lepidoptera) in the genera *Agassiziella, Elophila* and *Parapoynx*, were not included in the analysis because these were reported only in indoor habitats (Roques et al., 2016), and consequently we did not consider them as established in the wild. Another aspect of the life cycle of freshwater insects that may impede invasions is their lack of adaptations amenable to long distance transport via shipping (e.g., with ballast water; Duggan et al. 2006; Karatayev et al., 2009; Liebhold et al., 2016). Aquatic invertebrates with numerous non-native species, such as crustaceans and molluscs, often have life stages that are adapted to transport with ballast water (*e*.*g*., durable eggs or planktonic larval stages or drought-resistance in adult stages), a trait known to facilitate invasions (Panov et al., 2004; Ricciardi, 2015). Most freshwater insects lack adaptations that would enable them to tolerate conditions that prevail in ballast water, such as low oxygen levels (Fenoglio et al., 2016). However, some representatives of the Diptera, such as mosquitoes (Culicidae) are well adapted to low oxygen conditions and consequently are easily transported and can survive even in small amounts of water (Benedict et al., 2007; Ibañez-Justicia, 2020; Medlock et al., 2012). Hitchhiking at the egg stage in substrates without biological significance such as used tyres and ornamental plants (*e*.*g*., “lucky bamboo” *Dracaena* spp.) is a common pattern favouring invasion in dipteran species such as the tiger mosquito, *Aedes albopictus* (Linthicum et al., 2003; Rabitsch, 2010).

### Life cycle adaptations and habitat requirements

Differences in life cycle adaptations and habitat requirements facilitating invasions are also likely to be involved in the lower invasion success of freshwater insects. Asexual reproduction such as parthenogesis is very common among terrestrial non-native insects (and freshwater invaders such as mollusks) (Brockerhoff & Liebhold, 2017; Karatayev et al., 2009; Lee et al., 2005; Peacock & Worner, 2008), but it appears to be generally rare in freshwater insects (de Moor, 1992; Fenoglio et al., 2016). However, asexual reproduction occurs in some freshwater insects such as certain Odonata (e.g., de Moor, 1992; Lorenzo-Carballa et al., 2011; 2012), one of few freshwater orders that includes successful invaders (Table S2), as well as in some mayflies (Liegeois et al., 2021).

Because most freshwater insects have an aquatic and a terrestrial stage, they require suitable freshwater and terrestrial habitat, which may be an impediment to successful establishment of non-native species. Many terrestrial non-native insects are disturbance-adapted and benefit from anthropogenic habitat modification (*e*.*g*., urbanisation or agriculture) and habitat disturbance which provide conditions that facilitate their establishment and spread (Liebhold et al., 2016; Lozon & MacIsaac, 1997). For example, disturbance-adapted terrestrial invaders are common in the orders Blattodea, Hemiptera, and Phthiraptera (Liebhold et al., 2016; Peck & Roth, 1992), which are over-represented orders (see Fig. 1). By contrast, freshwater insects typically prefer undisturbed habitats (Rosenberg & Resh, 1993), and, consequently, they are less likely to benefit from anthropogenic habitat alteration. This is partly explained by the lack of tolerance of low-oxygen conditions which makes survival and establishment in disturbed, modified or polluted aquatic habitats, where propagules typically arrive, less likely. We found that the primarily aquatic orders Ephemeroptera, Plecoptera and Trichoptera (EPT), most of which are sensitive to deterioration of water quality (Ab Hamid & Md Rawi, 2016; Hering et al., 2004; Rosenberg & Resh, 1993), are consistently under-represented. By contrast, most established non-native freshwater invertebrates show some tolerance to organic pollutants and low dissolved oxygen (Karatayev et al., 2009). Unlike the EPT, some mosquitoes (Culicidae), hoverflies (Syrphidae) and some other Diptera that are adapted to poor water quality and low oxygen conditions can survive and become established even in small artificial aquatic habitats (Benedict et al., 2007; Derraik, 2005).

The ecological niche of many herbivorous terrestrial invasive insects is determined by the occurrence of their host plants or close relatives, although shifts to novel plants are predictable to some degree (Mech et al., 2019; Pearse & Altermatt, 2013). However, herbivory on vascular plants is considered a less important feeding category in freshwater insects where filter-feeding on phytoplankton or detritus dominate (Allan et al., 2020). Vascular plants are not dominant in most freshwater systems, which probably contributes to the limited number of non-native freshwater insects (Pearse & Altermatt, 2013).

Apart from the Diptera, Coleoptera and Hemiptera also have many representatives that are successful invaders (de Moor, 1992; Liebhold et al., 2021). However, these orders are among the biggest and most diverse insect orders (*e*.*g*., Skevington & Dang, 2002; Stork, 2018), and for such diverse groups it is difficult to determine a single mechanism underpinning the success of invasions. In these cases, an analysis on a scale of families or genera, as conducted for Coleoptera by Liebhold et al. (2021), may more accurately address mechanisms of invasion.

Using either the strict or the liberal dataset (based on strict or liberal definitions of “aquatic” insect - see methods) did not affect the overall results. The only difference occurred in the significance levels of the comparison between the strict and liberal datasets for New Zealand where the difference in the ratios of non-native to native species between freshwater and terrestrial habitats in the strict dataset was marginally non-significant. This is probably due to the limited richness range of native freshwater species in New Zealand. As this difference is significant in the liberal dataset for New Zealand, and the results were otherwise similar between the strict and liberal datasets for Europe and North America, we conclude that the applied definition of freshwater species does not generally affect the overall results. However, the definition of ‘aquatic species’ differs across various sources (compare: Karatayev et al., 2009; Merritt & Cummins, 1996; Merrit et al., 2008), which introduces uncertainty in comparisons among studies and the conclusions that can be drawn. Interestingly, we observed that some non-native freshwater species are herbivores feeding on aquatic host plants (Jäch & Balke, 2008; Mor et al., 2010). As these species can be considered semi-aquatic, we included them in the liberal dataset. They included representatives of Coleoptera, Diptera and Lepidoptera, orders which were noticeably more species-rich in the liberal dataset, although they remained under-represented.

## 5. Conclusions

To conclude, we provide broad and consistent evidence that invasions of freshwater insect species are relatively rare, in contrast to terrestrial insects which are particularly well represented among invasive species. We show this pattern to be repeated across three world regions (Europe, North America and New Zealand). Two non-exclusive causes are likely to be responsible for this difference: (i) transport pathways facilitating invasions (*i*.*e*., international trade) are less effective in moving freshwater insects than terrestrial insects, and (ii) characteristics of the life cycles and habitat requirements of freshwater insects predispose them to be less invasive than terrestrial insects. The alternative hypothesis, namely that the differential invasion success is due to freshwater habitats being less invasible than terrestrial habitats, is less plausible, because freshwater habitats are actually highly invaded by other macro-invertebrates (e.g., molluscs, crustaceans) (Karatayev et al. 2009; Ricciardi, 2015). We highlighted several more detailed mechanisms that are likely to contribute to the apparent causes (i) and (ii) given above. However, a more thorough understanding of these mechanisms would require more comprehensive experimental approaches and finer taxonomic resolution. Furthermore, understanding these mechanisms can improve predictions of future invaders and their impacts (Pyšek et al., 2012) and provide insights into broader ecological and evolutionary processes (Sax et al., 2007) that play a role in invasions. Future studies should aim at explaining, for example, the invasion patterns observed in diverse groups such as Coleoptera or Diptera. Of considerable importance would also be an improved understanding of the implications of potential differential effects of climate change and other global change drivers on invasions across freshwater and terrestrial habitats.

## Acknowledgements

We thank the ETH Board for funding through the Blue-Green Biodiversity (BGB) Initiative (BGB2020). AML acknowledges support from USDA Forest Service International Programs and grant EVA4.0, No. CZ.02.1.01/0.0/0.0/16_019/0000803 financed by Czech Operational Programme Science.

## Data accessibility statement

All relevant data is contained in the manuscript’s supplementary files.

