## Supplemental text and tables Table S3, Table S4 for "Fewer non-native insects in freshwater than in terrestrial habitats across continents"

**Supplementary materials**

**Table S1.** Number of aquatic and terrestrial insect species across insect orders in Europe, North America and New Zealand. This data used for the analysis.

(Attached as a separate csv file)

**Table S2.** List of non-native freshwater insect species established in Europe, New Zealand and North America.

(Attached as a separate MS Excel file).

**Table S3.** Details on the single term deletion process, applied to obtain the most parsimonious models on the basis of the Χ^2^ test. Statisically significant *p* values are highlighted in bold.

| **Strict dataset** | | | | |
| --- | --- | --- | --- | --- |
| *Terms* | *D.f.* | *Deviance* | *Scaled deviance* | *p* |
| Native*Habitat | 1 | 842.00 | 2.58 | 0.1082 |
| Native*Region | 2 | 895.45 | 8.28 | **0.0159** |
| Habitat*Region | 2 | 827.65 | 1.05 | 0.59093 |
| **Liberal dataset** | | | | |
| *Terms* | *D.f.* | *Deviance* | *Scaled deviance* | *p* |
| Native*Habitat | 1 | 880.27 | 1.25 | 0.2625 |
| Native*Region | 2 | 955.53 | 8.94 | **0.0114** |
| Habitat*Region | 2 | 869.32 | 0.13 | 0.9342 |
| *D.f.* - degrees of freedom, *p* - p value obtained from the Χ^2^test | | | | |

**Table S4.** Pairwise comparisons of the marginal means obtained from the binomial regression analysis of the strict and liberal datasets. Presented p values are adjusted by the Benjmaini & Hochberg correction for three comparisons.

| **Strict dataset** |  |  |  |  |
| --- | --- | --- | --- | --- |
| *Comparison* | *Estimates* | *S.E.* | *t.ratio* | *p value* |
| freshwater Europe-terrestrial Europe | -0.468 | 0.1041 | -4.498 | <0.0001 |
| Freshwater New Zealand-terrestrial New Zealand | -0.174 | 0.1020 | -1.711 | 0.0905 |
| Freshwater North American-terrestrial North America | -0.569 | 0.0597 | -9.530 | <0.0001 |
| **Liberal dataset** | | | | |
| *Comparison* | *Estimates* | *S.E.* | *t.ratio* | *p value* |
| Freshwater Europe-terrestrial Europe | -0.300 | 0.0737 | -4.073 | 0.0001 |
| Freshwater New Zealand-terrestrial New Zealand | -0.391 | 0.0832 | -4.701 | <0.0001 |
| Freshwater North American-terrestrial North America | -0.498 | 0.0457 | -10.881 | <0.0001 |

**Note S1.** Details on data compilation.

When data on the number of species for a given group was not directly available, we determined the number by subtracting it from the total number of species in a given order (Table 2). For instance, the number of native terrestrial insect species in Europe was calculated as the difference between the total number of native insect species in Europe (obtained from de Jong *et al.* 2014) and the number of freshwater species in Europe (obtained from the freshwaterecology.info database, Schmidt-Kloiber & Hering 2015). In the case of non-native species of North America, we reviewed the list of non-native species, selecting those which are aquatic according to the applied definition.

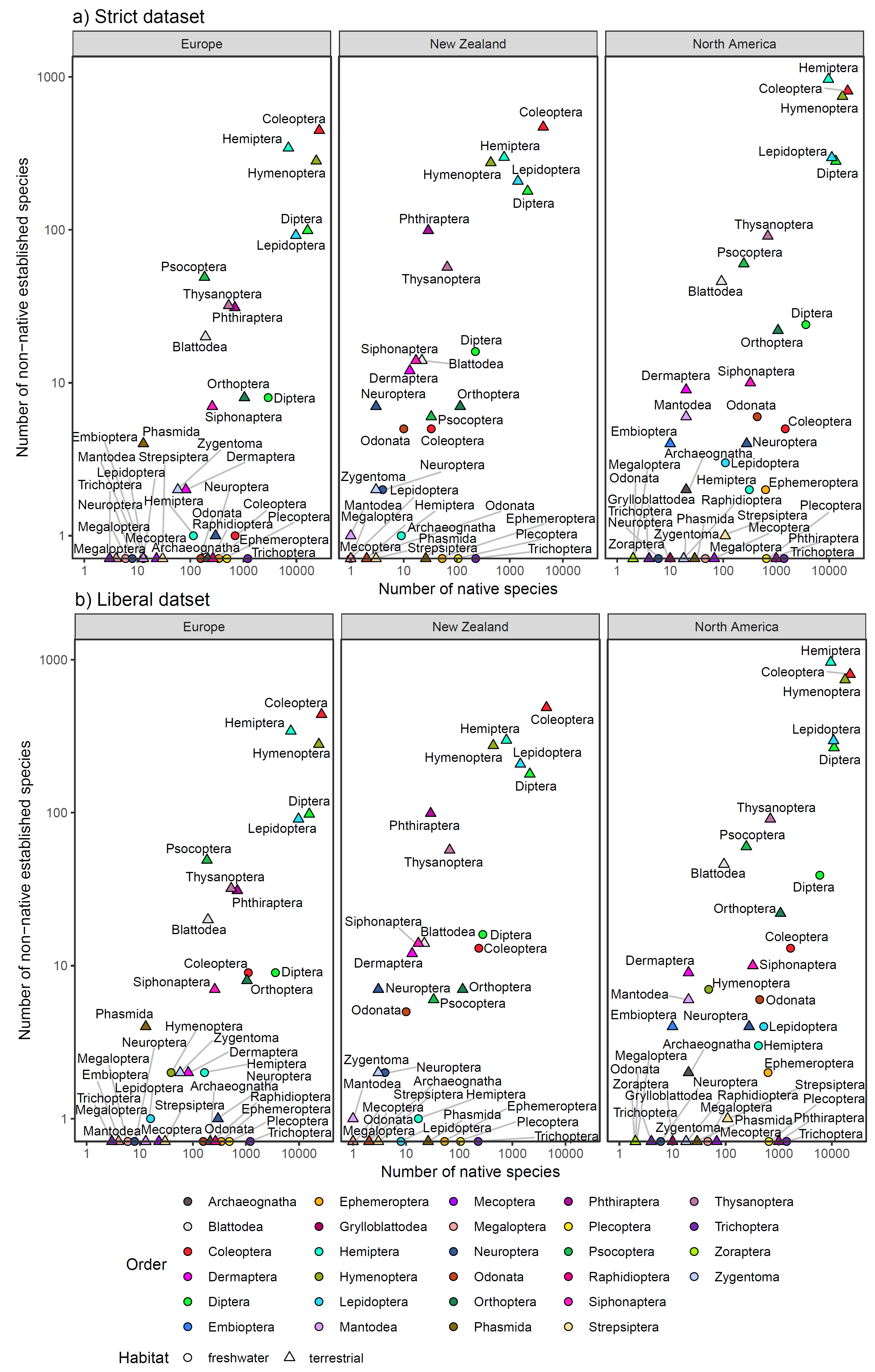

**Fig. S1.** Numbers of non-native and native species in freshwater and terrestrial insect orders, across the three the investigated regions (Europe, New Zealand, North America). Different orders are highlighted by colours.

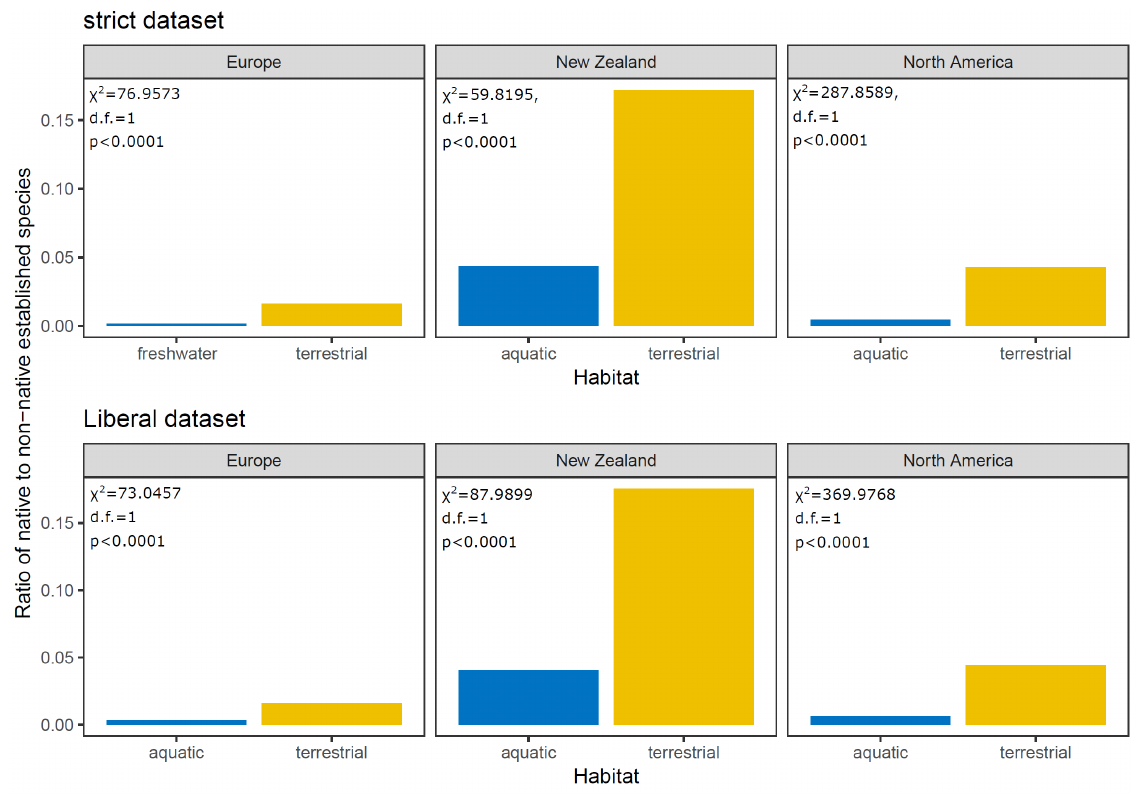
**Fig. S2.** Ratio of non-native to native species in freshwater and terrestrial habitats. Presented values correspond to X^2^ test of independence, calculated on the full species sample of freshwater and terrestrial species, for each of the regions.
