## Supplementary figures and images for "Fewer non-native insects in freshwater than in terrestrial habitats across continents"

### Supplemental figure: Fig. S1

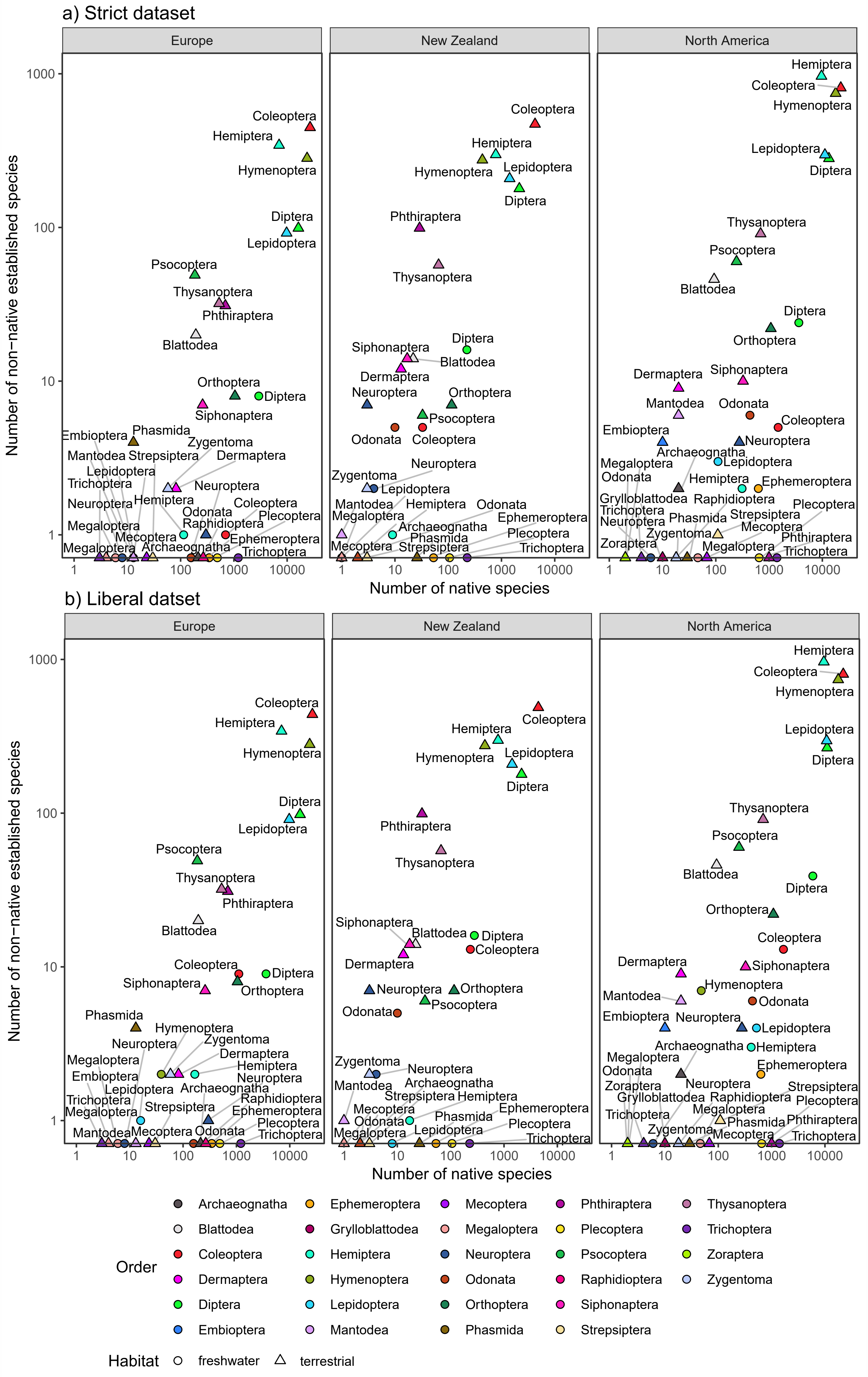

### Supplemental figure: Fig. S2

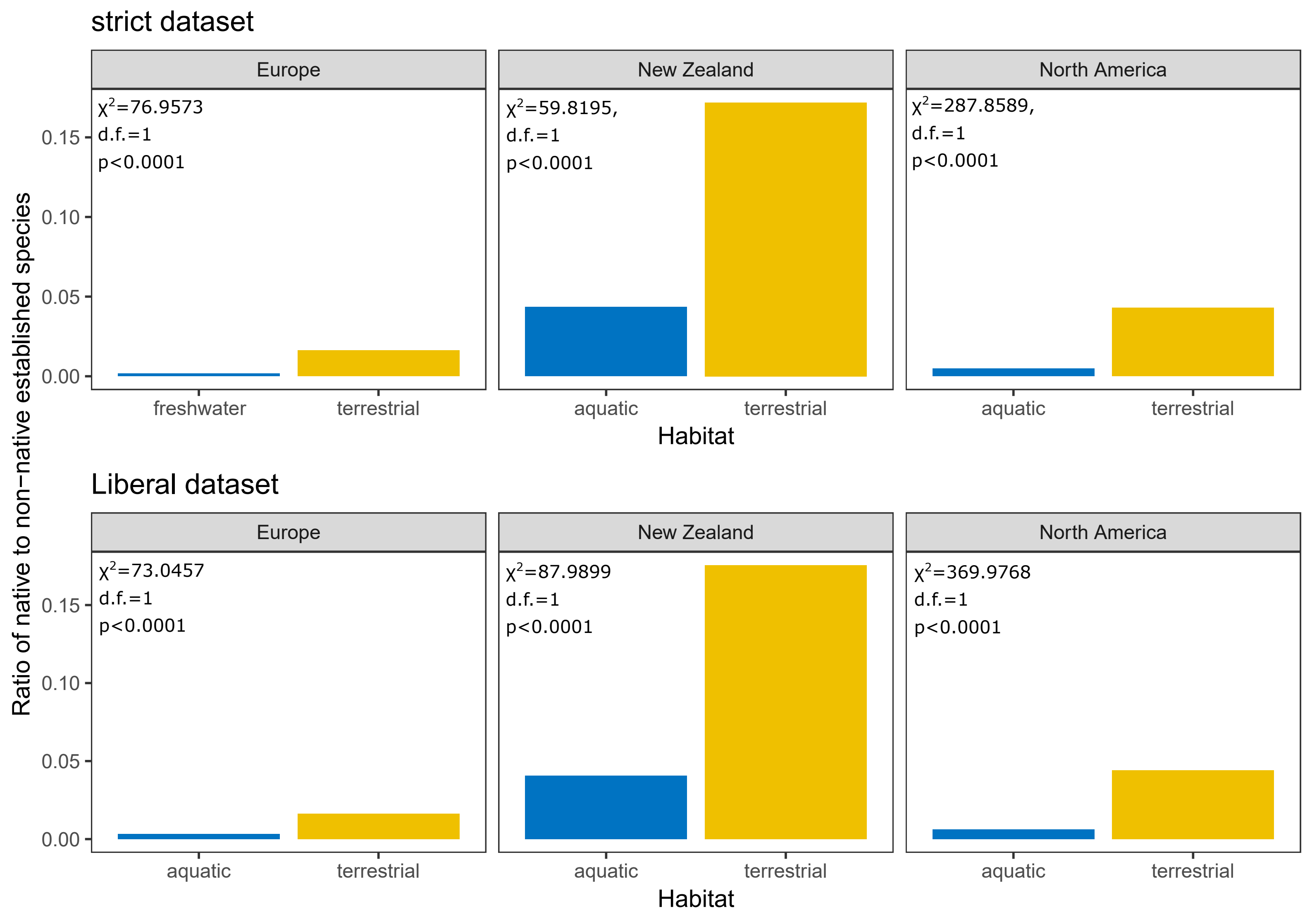
